# Mutant GT198 in angiogenesis as a common origin of human prostate and bladder cancers

**DOI:** 10.1101/726679

**Authors:** Liyong Zhang, Yehai Liu, Liang Cheng, Chengquan Zhao, Lan Ko

**Affiliations:** Georgia Cancer Center, Department of Pathology, Medical College of Georgia, Augusta University, Augusta, GA, USA; Department of Otolaryngology-Head & Neck Surgery, The First Affiliated Hospital, Anhui Medical University, Hefei, Anhui, China; Department of Pathology and Laboratory Medicine, Indiana University School of Medicine, Indianapolis, IN, USA; Department of Pathology, Magee-Womens Hospital, University of Pittsburgh, Pittsburgh, PA, USA

**Keywords:** Oncogene GT198, somatic mutations, tumor stroma, prostate and bladder cancer

## Abstract

Prostate and bladder cancers are common cancers in men. It has been speculated that the high concomitant incidence of the two cancers is due to a potential shared cause underlying both cancers. In this report, we have identified a common cause of human prostate and bladder cancers as the mutant oncoprotein GT198 (*PSMC3IP*). GT198 is a DNA repair factor and a steroid hormone receptor coactivator. GT198 has been previously shown to be mutated in angiogenic pericyte stem cells in solid tumor microenvironment. GT198 is also a direct protein target of chemo drugs paclitaxel and doxorubicin. Here we show, the *GT198* gene is mutated with protein overexpression in tumor stroma of human prostate and bladder cancers. Affected stromal cells include angiogenic blood vessel pericyte stem cells, and vascular smooth muscle cell lineages including myofibroblasts in prostate and smooth muscle cells in bladder. In prostate cancers, GT198^+^ tumor stromal cells are associated with early stages of cancer with lower Gleason scores. In bladder cancers, the presence of angiogenesis and GT198^+^ stroma are associated with better progression-free survival in docetaxel-treated patients. Together, our evidence suggests that angiogenic pericyte stem cells are initial lesions producing a mutant stroma carrying *GT198* somatic mutations. Subsequently, mutant myofibroblasts promote adenocarcinomas in prostate and mutant smooth muscle cells promote urothelial carcinomas in bladder. Chemo drugs targeting to GT198 is more effective in early stages of cancers with GT198^+^ stromal cells. This study supports oncoprotein GT198 as a common cause and a drug target in human prostate and bladder cancers.

## INTRODUCTION

Prostate cancer and bladder cancer are prevalent cancers in men. Clinical evidence has previously suggested that prostate and bladder cancers have higher concomitant incidence with a potential common pathway in tumor development (1–3).

In this study, we propose that a common molecular lesion exists as a shared origin of human prostate and bladder cancers. The responsible oncogene is called *GT198* (gene symbol *PSMC3IP*), its protein product is also called Hop2 and TBPIP.

GT198 was initially identified as a transcriptional coactivator stimulating steroid hormone receptor-mediated gene activation (4,5). GT198 directly binds to androgen, estrogen, and progesterone receptors (4), and stimulates CYP17 (cytochrome P450 17α) promoter (6). GT198 is potentially a master regulator in steroid hormone biosynthesis, which explains its oncogenic involvement in human breast (7), ovarian (6), and fallopian tube cancers (8). Prostate and bladder cancers are also hormone-dependent (9–13).

GT198 is also a DNA repair factor known as mammalian Hop2 or TBPIP (8,14,15). GT198 is a DNA-binding protein dimer, binds to single- and double-stranded DNAs. It has been extensively characterized in DNA repair by participating in homologous DNA recombination (16–19). Since DNA repair factors are working as a protein complex, it should be no surprising for the extensive evidence that DNA repair factors including BRCA1 and BRCA2 contribute to the risk of prostate and bladder cancers (20), because GT198 is one of them in the DNA repair machinery.

It is important to note that the human *GT198* gene is located at the chromosome 17q21 locus, 470 Kb proximal to the *BRCA1* gene (21,22). The 17q21 locus has been repeatedly identified as a prostate cancer susceptibility gene locus by previous cancer genetic studies in familial prostate cancers (23–27). One early study correctly narrowed down the prostate cancer candidate gene to a less than 2 Mb genomic interval between polymorphic marker *D17S776* and *BRCA1* (28), where *GT198* is located. Evidence for the involvement of DNA repair genes in inherited prostate cancer is also significant (29,30). Despite mounting evidence, *GT198* was not revealed by cancer genetic studies alone in part due to its very small gene size, shadowed by *BRCA1*, and carrying mostly splicing mutations rather than truncation mutations (see below).

The human *GT198* gene has previously been shown to link to breast and ovarian cancer predisposition. Germline mutations in *GT198* are identified in familial and early-onset human breast and ovarian cancer (21,31), and in familial ovarian disease (32). Somatic mutations in *GT198* are frequent and recurrent in sporadic breast (7), and ovarian cancers (6), where mutant GT198 protein leads to constitutive transcriptional activation (8,21). In human breast cancer, the *GT198* gene is mutated in the tumor stromal microenvironment. The mutant cells are pericyte stem cells and the descendent vascular smooth muscle cell lineage, including myoepithelial cells, fibroblasts, and adipocytes (7). Consistent with our findings, breast and prostate cancers share profound similarities in numerous aspects of their pathophysiology (12). Myoepithelial cells in breast are functionally equivalent to myofibroblasts in prostate. Both are contractile, hormone-responsive, and belong to the vascular smooth muscle cell lineage. When blood vessel pericytes were mutated in *GT198* during angiogenesis (6,7,33), stroma would become mutated consisting mutant stromal cells. Extensive evidence have already supported the important roles of stromal cells (34,35), prostate cancer-associated fibroblasts (CAF) (36), and myofibroblasts (37), in tumorigenesis. Except that tumor stromal cells should no longer be considered normal, they are malignant carrying mutations prior to the alteration of epithelial cells. This may be common in human solid tumors (38), where pericyte stem cells are affected by overexpressing GT198 (33).

Taxanes including paclitaxel and docetaxel are chemotherapy drugs for prostate cancer. It is urgent to identify new therapeutic biomarkers in responders for effective treatment. Paclitaxel is lately found to be a direct GT198 inhibitor (38,39). Thus, GT198 is a previously unanticipated target of taxane and needs to be evaluated as a new biomarker of taxane responsiveness in prostate cancer treatment.

In this report, we found that the *GT198* gene is mutated with protein overexpression in human prostate and bladder cancer stroma. The earliest changes are angiogenic blood vessels, which consist GT198^+^ pericyte stem cells. Extensive stromal proliferation is found to follow angiogenesis. Myofibroblasts in prostate cancer and smooth muscle cells in bladder cancer are the most populous GT198^+^ stromal cells, and carry somatic mutations in *GT198*. In prostate cancers, GT198^+^ stromal cells are associated with early cancers with lower Gleason scores. In bladder cancers, GT198^+^ stroma but not GT198^+^ tumor is associated with better progression-free survival in docetaxel-treated patients. Our data together suggest that mutant GT198^+^ stromal cells are initial lesions in both prostate and bladder cancers before epithelial tumor cell development. Docetaxel targeting to GT198^+^ stroma with angiogenesis may result in higher treatment efficacy. This is a first study to support oncoprotein GT198 as a common cause as well as a treatment biomarker in human prostate and bladder cancers.

## RESULTS

### GT198 overexpression in human prostate and bladder tumor stroma

We have previously identified cytoplasmic GT198 expression as precursor lesions in tumor stroma of various human solid tumors (6,7,33). In this study, focusing on GT198 in human prostate and bladder cancers, we analyzed GT198 expression by immunohistochemistry in 65 cases of tumor tissue microarrays and 15 cases of tumor sections of prostate cancers, and 17 cases of tumor sections of bladder cancers (**Table 1–4**).

**Table 1.** GT198 expression in prostate tumor tissue microarray (Novus Biologicals NBP2-30169, IMH-303) Overall GT198 positive rate = 97.5% (39/40) Two prostate tumor tissue microarrays (40 cases in NBP2-30169/IMH-303, core diameter 2.0 mm, thickness 4 μM; and 25 cases in PR1001, core diameter 1.0 mm, thickness 5 μM.) were immunohistochemically stained with GT198 and analyzed for the presence of GT198^+^ myofibroblasts and GT198^+^ tumor cells. The GT198 negative cases are shaded in light blue. Percent of positive cases in total and percent of positive cases in myofibroblasts or tumor cells are listed in blue. In NBP2-30169, there are nine cases of matched adjacent tissues absent of tumor cells but containing positive tumor stroma. The quality of the NBP2-30169 section is much higher than the PR1001, in which immunohistochemical staining may be failed in some negative cases.

| Core No | Age | Diagnosis | Gleason Score | pTNM | Stage | GT198+ Myofibroblast | GT198+ Tumor |
| --- | --- | --- | --- | --- | --- | --- | --- |
| 1 | 60 | adenocarcinoma | 9 | T3bN0M0 | III |  | + |
| 2 | 64 | adenocarcinoma | 7 | T2cN0M0 | IIB | + |  |
| 3 | 71 | adenocarcinoma | 9 | T3bN0M0 | III |  | + |
| 4 | 64 | adenocarcinoma | 10 | T3aN0M0 | III | + | + |
| 5 | 59 | adenocarcinoma | 9 | T3bN0M0 | III | + |  |
| 6 | 65 | adenocarcinoma | 8 | T3bN0M0 | III |  | + |
| 7 | 73 | adenocarcinoma | 7 | T2cN0M0 | IIB | + |  |
| 8 | 69 | adenocarcinoma | 7 | T2cN0M0 | IIB | + |  |
| 9 | 62 | adenocarcinoma | 7 | T2cN0M0 | IIB | + |  |
| 10 | 66 | adenocarcinoma | 9 | T3bN0M0 | III |  | + |
| 11 | 60 | adenocarcinoma | 9 | T3bN0M0 | III | + |  |
| 12 | 66 | adenocarcinoma | 7 | T3aN0M0 | III | + |  |
| 13 | 70 | adenocarcinoma | 7 | T3bN0M0 | III | + |  |
| 14 | 65 | adenocarcinoma | 9 | T3bN0M0 | III | + |  |
| 15 | 67 | adenocarcinoma | 9 | T3bN1M0 | III |  | + |
| 16 | 69 | adenocarcinoma | 7 | T3bN0M0 | III | + |  |
| 17 | 63 | adenocarcinoma | 9 | T3aN1M0 | III |  | + |
| 18 | 69 | adenocarcinoma | 7 | T3aN0M0 | III | + |  |
| 19 | 70 | adenocarcinoma | 7 | T3aN0M0 | III | + |  |
| 20 | 58 | adenocarcinoma | 9 | T3bN0M0 | III | + | + |
| 21 | 58 | adenocarcinoma | 7 | T3bN0M0 | III | + |  |
| 22 | 71 | adenocarcinoma | 7 | T2cN0M0 | IIB | + |  |
| 23 | 70 | adenocarcinoma | 7 | T3bN0M0 | III | + | + |
| 24 | 59 | adenocarcinoma | 6 | T2bN0M0 | IIA | + |  |
| 25 | 63 | adenocarcinoma | 9 | T3bN0M0 | III | + |  |
| 26 | 72 | adenocarcinoma | 9 | T3bN0M0 | III |  | + |
| 27 | 66 | adenocarcinoma | 8 | T3bN0M0 | III | + |  |
| 28 | 70 | adenocarcinoma | 6 | T3bN0M0 | III |  |  |
| 29 | 70 | adenocarcinoma | 7 | T2cN0M0 | IIB | + | + |
| 30 | 68 | adenocarcinoma | 8 | T3bN0M0 | III | + |  |
| 31 | 63 | adenocarcinoma | 10 | T3bN0M0 | III |  | + |
| 32 | 57 | adenocarcinoma | 7 | T3bN0M0 | III | + |  |
| 33 | 72 | adenocarcinoma | 8 | T2cN0M0 | IIB |  | + |
| 34 | 70 | adenocarcinoma | 8 | T3bN0M0 | III | + |  |
| 35 | 75 | adenocarcinoma | 9 | T3bN0M0 | III |  | + |
| 36 | 62 | adenocarcinoma | 9 | T3bN0M0 | III |  | + |
| 37 | 63 | adenocarcinoma | 9 | T3bN0M0 | III |  | + |
| 38 | 53 | adenocarcinoma | 9 | T3bN0M0 | III |  | + |
| 39 | 63 | adenocarcinoma | 8 | T3bN0M0 | III | + | + |
| 40 | 44 | adenocarcinoma | 7 | T3bN0M0 | III | + |  |
|  |  |  |  |  |  | **65%** | **45%** |
| 41 | 69 | adjacent (match of #8) | . | . | . | + |  |
| 42 | 62 | adjacent (match of #9) | . | . | . | + |  |
| 43 | 66 | adjacent (match of #12) | . | . | . | + |  |
| 44 | 65 | adjacent (match of #14) | . | . | . | + |  |
| 45 | 69 | adjacent (match of #18) | . | . | . | + |  |
| 46 | 70 | adjacent (match of #19) | . | . | . | + |  |
| 47 | 70 | adjacent (match of #29) | . | . | . | + |  |
| 48 | 63 | adjacent (match of #31) | . | . | . | + |  |
| 49 | 44 | adjacent (match of #40) | . | . | . | + |  |
|  |  |  |  |  |  | **100%** | **0%** |

**Table 2.** GT198 expression in prostate tumor tissue microarray (US Biomax PR1001) Overall GT198 positive rate = 68% (17/25) Two prostate tumor tissue microarrays (40 cases in NBP2-30169/IMH-303, core diameter 2.0 mm, thickness 4 μM; and 25 cases in PR1001, core diameter 1.0 mm, thickness 5 μM.) were immunohistochemically stained with GT198 and analyzed for the presence of GT198^+^ myofibroblasts and GT198^+^ tumor cells. The GT198 negative cases are shaded in light blue. Percent of positive cases in total and percent of positive cases in myofibroblasts or tumor cells are listed in blue. In NBP2-30169, there are nine cases of matched adjacent tissues absent of tumor cells but containing positive tumor stroma. The quality of the NBP2-30169 section is much higher than the PR1001, in which immunohistochemical staining may be failed in some negative cases.

| **Table 2. GT198 expression in prostate tumor tissue microarray (US Biomax PR1001)**<br>**Overall GT198 positive rate = 68% (17/25)** |  |  |  |  |  |  |  |
| --- | --- | --- | --- | --- | --- | --- | --- |
| Core No | Age | Diagnosis | Gleason Score | pTNM | Stage | GT198+ Myofibroblast | GT198+ Tumor |
| 1 | 70 | adenocarcinoma | 7 | T3N0M0 | III |  | + |
| 2 | 64 | adenocarcinoma | 6 | T2aN0M0 | II | + |  |
| 3 | 78 | adenocarcinoma | 7 | T4N1M1b | IV |  |  |
| 4 | 65 | adenocarcinoma | 8 | T2N0M0 | II | + |  |
| 5 | 75 | adenocarcinoma | 6 | T4N1M1 | IV |  |  |
| 6 | 70 | adenocarcinoma | 7 | T2N1M1c | IV |  |  |
| 7 | 62 | adenocarcinoma | 7 | T3N1M1h | IV |  |  |
| 8 | 76 | adenocarcinoma | 7 | T2aN0M0 | II | + |  |
| 9 | 62 | adenocarcinoma | 10 | T2N0M0 | II | + |  |
| 10 | 64 | adenocarcinoma | 9 | T2N0M0 | II | + |  |
| 11 | 65 | adenocarcinoma | 7 | T2N0M0 | II |  | + |
| 12 | 82 | adenocarcinoma | 10 | T3N2M1 | IV | + |  |
| 13 | 66 | adenocarcinoma | 7 | T3aN0M0 | III | + |  |
| 14 | 76 | adenocarcinoma | 10 | T3N1M1b | IV | + |  |
| 15 | 76 | adenocarcinoma | 7 | T2aN0M0 | II | + |  |
| 16 | 73 | adenocarcinoma | 10 | T3N1M1 | IV |  | + |
| 17 | 60 | adenocarcinoma | 7 | T3N1M0 | IV | + |  |
| 18 | 73 | adenocarcinoma | 10 | T4N1M1c | IV | + |  |
| 19 | 77 | adenocarcinoma | 7 | T2N0M0 | II | + |  |
| 20 | 55 | adenocarcinoma | 10 | T2N0M0 | II | + |  |
| 21 | 60 | adenocarcinoma | 10 | T2N0M0 | II |  |  |
| 22 | 87 | adenocarcinoma | 10 | T2N0M0 | II |  |  |
| 23 | 81 | adenocarcinoma | 9 | T2aN0M0 | III |  | + |
| 24 | 75 | adenocarcinoma | 10 | T2N1M1c | IV |  |  |
| 25 | 80 | adenocarcinoma | 6 | T4N1M1c | IV |  |  |
|  |  |  |  |  |  | **52%** | **16%** |

**Table 3.** GT198 expression in human prostate tumor sections. Fifteen cases of human prostate tumor tissue FFPE sections were immunohistochemically stained with GT198 and analyzed for the presence of GT198^+^ blood vessels, myofibroblasts, and tumor cells. *GT198* mutations were only analyzed in cases P1-P4. The GT198 negative case is shaded in light blue. Percent of positive cases in total, percent of positive cases in each cell types, and percent of mutation positive cases are listed in blue. n.a., not analyzed.

| **Table 3. GT198 expression in prostate tumor slides**<br>**Overall GT198 positive rate = 93.3% (14/15)** |  |  |  |  |  |  |  |
| --- | --- | --- | --- | --- | --- | --- | --- |
| Case No | Age | Diagnosis | Gleason Score | GT198+ Blood vessel | GT198+ Myofibroblast | GT198+ Tumor | GT198 Mutation |
| P1 | 80 | adenocarcinoma | 6 | + | + | + | 6 |
| P2 | 85 | adenocarcinoma | 8 |  | + | + | 5 |
| P3 | 92 | adenocarcinoma | 9 | + | + |  | 1 |
| P4 | 90 | adenocarcinoma | 8 | + |  |  | 0 |
| P5 | 57 | adenocarcinoma | 8 | + | + | + | n.a. |
| P6 | 60 | adenocarcinoma | 7 | + | + | + | n.a. |
| P7 | 63 | adenocarcinoma | 7 | + | + | + | n.a. |
| P8 | 63 | adenocarcinoma | 8 | + | + | + | n.a. |
| P9 | 52 | adenocarcinoma | 7 | + | + | + | n.a. |
| P10 | 63 | adenocarcinoma | 9 |  |  |  | n.a. |
| P11 | 68 | adenocarcinoma | 7 | + | + |  | n.a. |
| P12 | 80 | adenocarcinoma | 7 | + | + | + | n.a. |
| P13 | 56 | adenocarcinoma | 7 |  | + |  | n.a. |
| P14 | 74 | adenocarcinoma | 6 | + | + | + | n.a. |
| P15 | 62 | adenocarcinoma | 8 | + | + |  | n.a. |
|  |  |  |  | 80.0% | 86.6% | 60% | 75% |

**Table 4.** GT198 expression in human bladder tumor sections. Seventeen cases of human bladder tumor FFPE sections with clinical information were immunohistochemically stained with GT198. The fifteen cases (B1-B15) were docetaxel-treated and were analyzed for the association of progression-free survival in **Figure 4**. The GT198 negative cases are shaded in light blue. Percent of positive cases in total, positive cases in cell types, angiogenesis, and mutation positive cases are listed in blue. pTNM, Pathological tumor-node-metastasis staging; PFS, Progression-free survival; n.a., not analyzed.

| **Table 4. GT198 expression in bladder tumor slides**<br>**Overall GT198 positive rate = 76.4% (13/17)** |  |  |  |  |  |  |  |  |  |  |  |  |
| --- | --- | --- | --- | --- | --- | --- | --- | --- | --- | --- | --- | --- |
| Case No | Age | Sex | pTNM | GT198+ Stroma | GT198+ Tumor | Angio-genesis | GT198 Mutation | Docetaxel Treated | PFS Month | Last Status | Progression Recurrence | Nodes Removed |
| B1 | 73 | F | pT0N0 | ++ |  | + | 2 | Yes | 33 | Alive | No | 0 of 9 |
| B2 | 72 | M | pT4aN0 | + | + |  | 0 | Yes | 19 | Dead | Liver, bone | 0 of 1 |
| B3 | 40 | F | pT3aNx |  | + |  | 1 | Yes | 12 | Dead | Peritoneum, lung | none |
| B4 | 44 | M | pT0N3 | + |  | + | 1 | Yes | 202 | Alive | No | 8 of 24 |
| B5 | 59 | M | pT3bN2 | + | + | + | 1 | Yes | 16 | Dead | Penis, scrotum | 4 of 4 |
| B6 | 61 | F | pT2aN2 |  |  |  | 0 | Yes | 4 | Dead | Liver, lung | 6 of 8 |
| B7 | 51 | M | pT0N0 | ++ |  | + | 0 | Yes | 182 | Alive | No | 0 of 11 |
| B8 | 51 | M | pT4aN1 |  | ++ |  | 0 | Yes | 9 | Dead | Pelvis | 4 of 11 |
| B9 | 61 | M | pT3aN1 | + | + |  | 0 | Yes | 138 | Alive | No | 1 of 15 |
| B10 | 53 | F | pT2bN1 | ++ |  | + | 4 | Yes | 23 | Dead | Liver | 1 of 3 |
| B11 | 63 | M | pT2bN0 |  |  |  | 0 | Yes | 5 | Dead | Unknown | 0 of 7 |
| B12 | 47 | M | pTisNx | + | + | + | 0 | Yes | 115 | Alive | No | none |
| B13 | 64 | F | pT3bN0 |  | ++ |  | 3 | Yes | 5 | Dead | Retroperitoneum | none |
| B14 | 70 | M | pT1N1 |  |  | + | 0 | Yes | 68 | Dead | Liver, lung | 1 of 5 |
| B15 | 57 | F | pT2aN1 |  |  |  | 0 | Yes | 45 | Dead | Groin nodes | 1 of 10 |
| B16 | 75 | M | pT3bN1 |  | + |  | n.a. | No | 19 | Dead | Unknown | 1 of 7 |
| B17 | 45 | M | pT2bN1 | + |  |  | n.a. | No | 132 | Alive | No | 1 of 14 |
|  |  |  |  | 52.9% | 47.0% | 41.1% | 40.0% |  |  |  |  |  |

T0: No evidence of a primary tumor. Ta: Noninvasive papillary carcinoma. Tis: Carcinoma *in situ* (CIS) or a flat tumor. T1: The tumor has spread to the connective tissue. T2: The tumor has spread to the muscle of the bladder wall. T3: The tumor has grown into the perivesical (fatty) tissue.
Figure credit: American Cancer Society

In human prostate stroma with angiogenesis before tumor cells appear, GT198 is strongly expressed in blood vessel pericytes in capillaries (**Figure 1A**). GT198 is also found in smooth muscle cells in larger blood vessels (**Figure 1B-C**), and in proliferative myofibroblasts of the prostate stroma at the early stages of cancer development (**Figure 1D**). GT198^+^ myofibroblasts continue to accumulate and become fibrous GT198^+^ tumor stroma (**Figure 1E**). More than half of the epithelial tumors are negative in GT198 expression, with diluted GT198^+^ myofibroblasts among them. Some evolve into GT198^+^ tumors (**Figure 1F-H**). Normal GT198^-^ blood vessels can also be found in tumor tissues.

**Figure 1.**
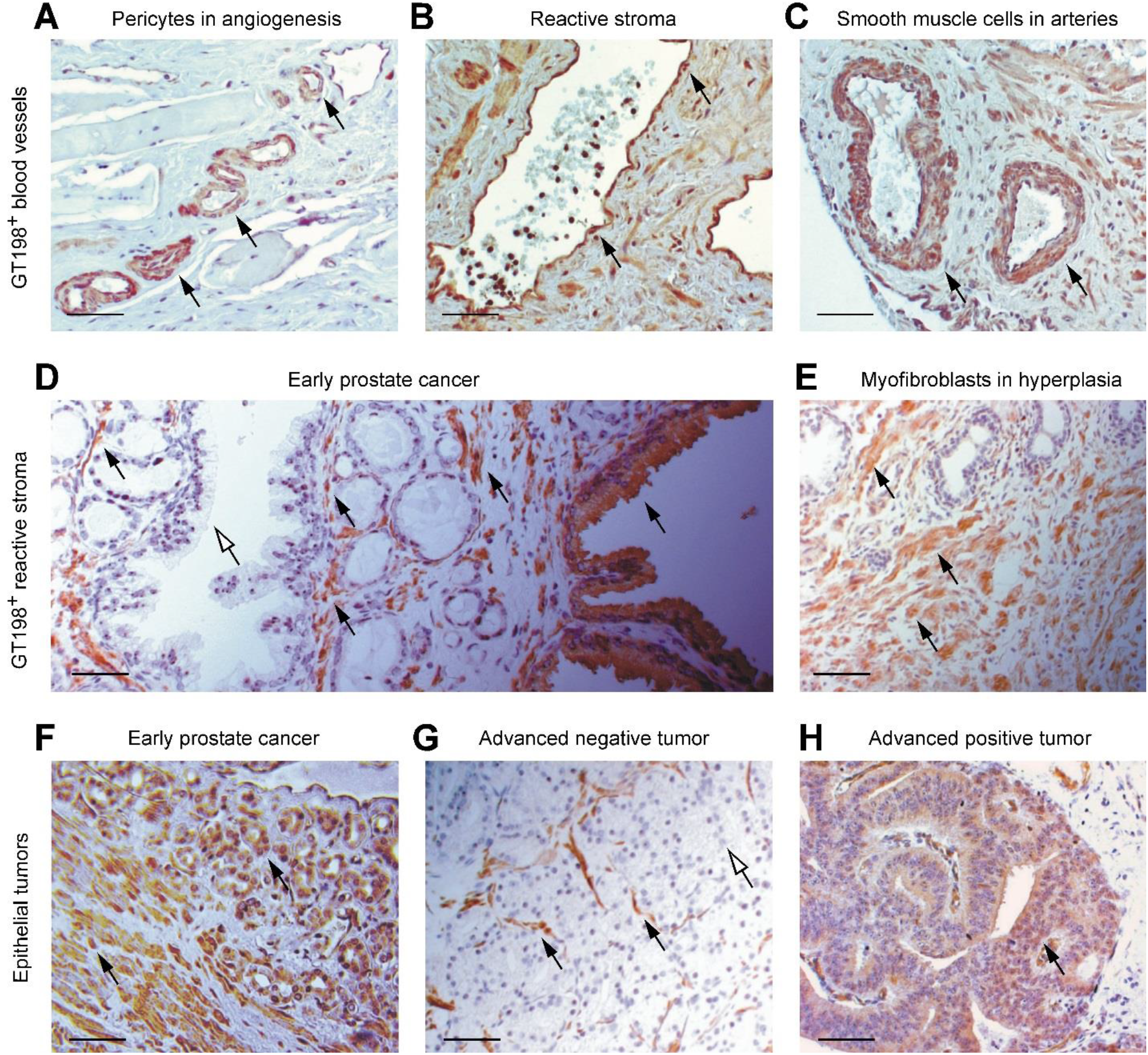
GT198 expression in human prostate cancers. Immunohistochemical staining of GT198. **(A)** At the early stage of prostate cancer development, GT198 is expressed in capillaries of tumor microenvironment with angiogenesis. Pericytes and vascular smooth muscle cells are GT198^+^. **(B-C)** In angiogenic prostate cancer stroma, both veins and arteries are GT198^+^ surrounded by proliferative fibroblasts and myofibroblasts. **(D)** In prostate cancers at the early stages, epithelium overgrowth is underlain by GT198^+^ myofibroblasts. Epithelial cells are often negative in GT198 expression except some clonal expanded clusters can be GT198^+^. **(E)** The proliferative and hyperplastic tumor stroma is populated by abundant GT198^+^ myofibroblasts. **(F)** Early stage GT198^+^ tumor cells and myofibroblasts. **(G-H)** In advanced prostate tumors, GT198^+^ myofibroblasts are diluted by the overgrowth of epithelial cells, either negative or positive in GT198 expression. Black and white arrows indicate GT198^+^ or GT198^-^ cells, respectively. Sections were counter-stained with hematoxylin. Scale bars = 100 μm.

With GT198 as a clear marker, it appears that GT198^+^ vessel pericytes as stem cells have produced GT198^+^ vascular smooth muscle cell lineages including myofibroblasts and blood vessel smooth muscles. These prostate stroma lesions together induce epithelial proliferation, given that myofibroblasts dictate prostate epithelial cell growth (13).

A very similar scenario can be found in human bladder cancers (**Figure 2**). In contrast to normal GT198^-^ blood vessels (**Figure 2A**), GT198^+^ angiogenic capillaries are clustered and underlie the urothelial layer (**Figure 2B**). The positive vessels further mature into larger vessels (**Figure 2C-D**), and the same vascular smooth muscle cell lineage eventually creates GT198^+^ smooth muscle layers of the bladder (**Figure 2E**). The fibrous GT198^+^ stroma can be predominant in the tissue slides, which permitted detection of *GT198* somatic mutations (see below). The stromal lesions are multifocal, derived from angiogenesis, and likely drive the growth of urothelial carcinomas (**Figure 2F**). Tumor cells can be either GT198^+^ or GT198^-^ (**Table 4**). It is currently unclear whether GT198^+^ stem or progenitor cells could differentiate into GT198^+^ urothelial carcinomas.

**Figure 2.**
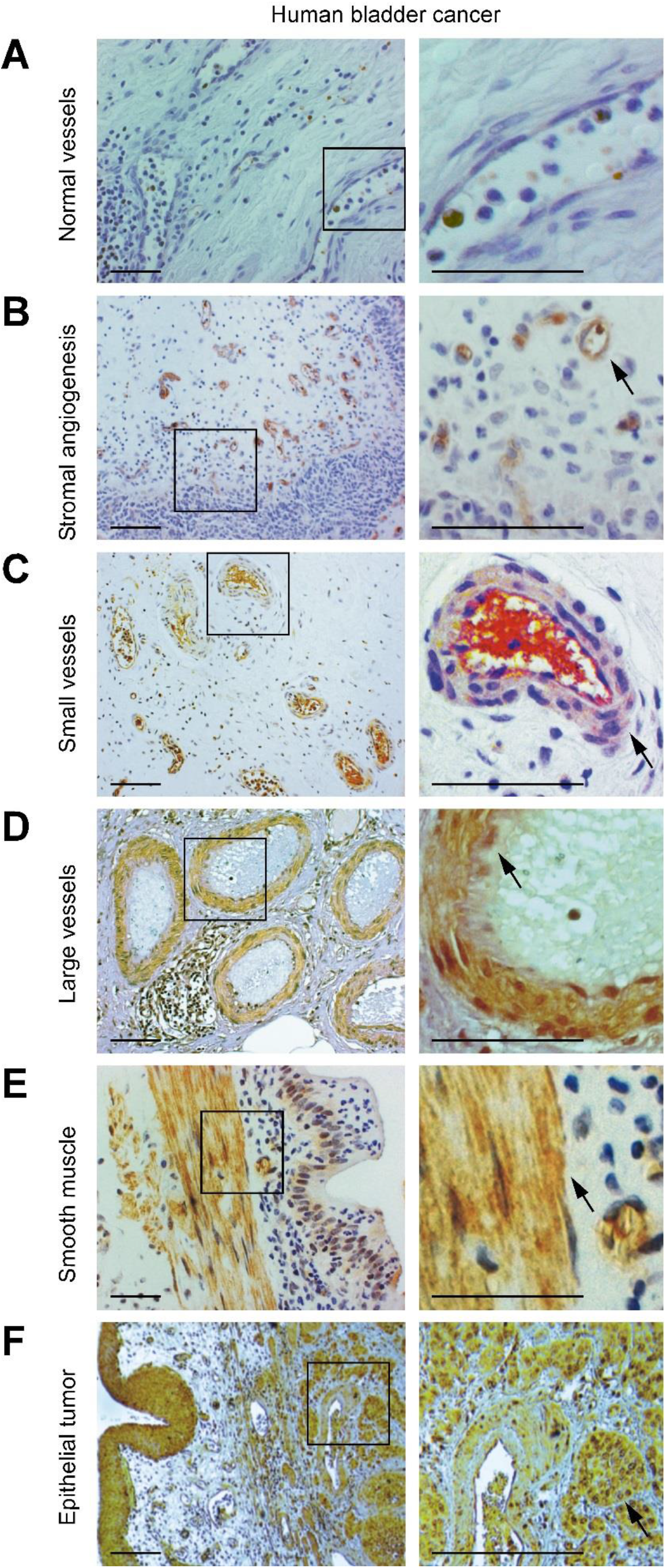
GT198 expression in human bladder cancers. Immunohistochemical staining of GT198. **(A)** An inflammatory bladder tumor stroma is served as a negative control in GT198 expression. Normal pericytes are devoid of GT198. A few white blood cells are GT198^+^. **(B)** GT198 is expressed in angiogenic blood vessels in stroma near the urothelial layer. Typically, abundant capillaries with surrounding fibroblasts are GT198^+^. **(C)** In angiogenic bladder cancer stroma, small blood vessels are GT198^+^. **(D)** In larger blood vessels, GT198 is robustly expressed in smooth muscle layers of the vessels. **(E)** In smooth muscles layers under the bladder urothelium, GT198 is strongly overexpressed. **(F)** In advanced bladder tumors, blood vessels and urothelial tumor cells are GT198^+^. Boxed areas are enlarged at right panels. Black arrows indicate GT198^+^ cells. Sections were counter-stained with hematoxylin. Scale bars = 100 μm.

### GT198^+^ stromal myofibroblasts are associate with low Gleason scores in prostate cancers

The expression of GT198 in prostate cancers was examined by immunohistochemical staining. The overall positive rates in prostate cancers are high, 97.5% (39/40) in a high quality tumor microarray, 68% (17/25) in a low quality microarray, and 93.3% (14/15) in prostate tumor sections (**Table 1–3**). In 9 cases of tumor adjacent stroma, traditionally called adjacent normal, GT198 is expressed in all cases (**Table 1** and **Figure 3A**). Because the size of tissue is much smaller in microarrays than tissue sections which contain more stromal areas, GT198^+^ myofibroblasts positive rate is found higher in tissue sections at 86.6% (**Figure 3A**). When Gleason scores were analyzed with the GT198 expression, GT198^+^ stroma is more associated with cases with low Gleason scores than the GT198^-^ stroma, but GT198^+^ tumors have no significant association with Gleason scores than the GT198^-^ tumors (**Figure 3C**). Data suggest that prostate cancers at the early stages contain more GT198^+^ myofibroblasts.

**Figure 3.**
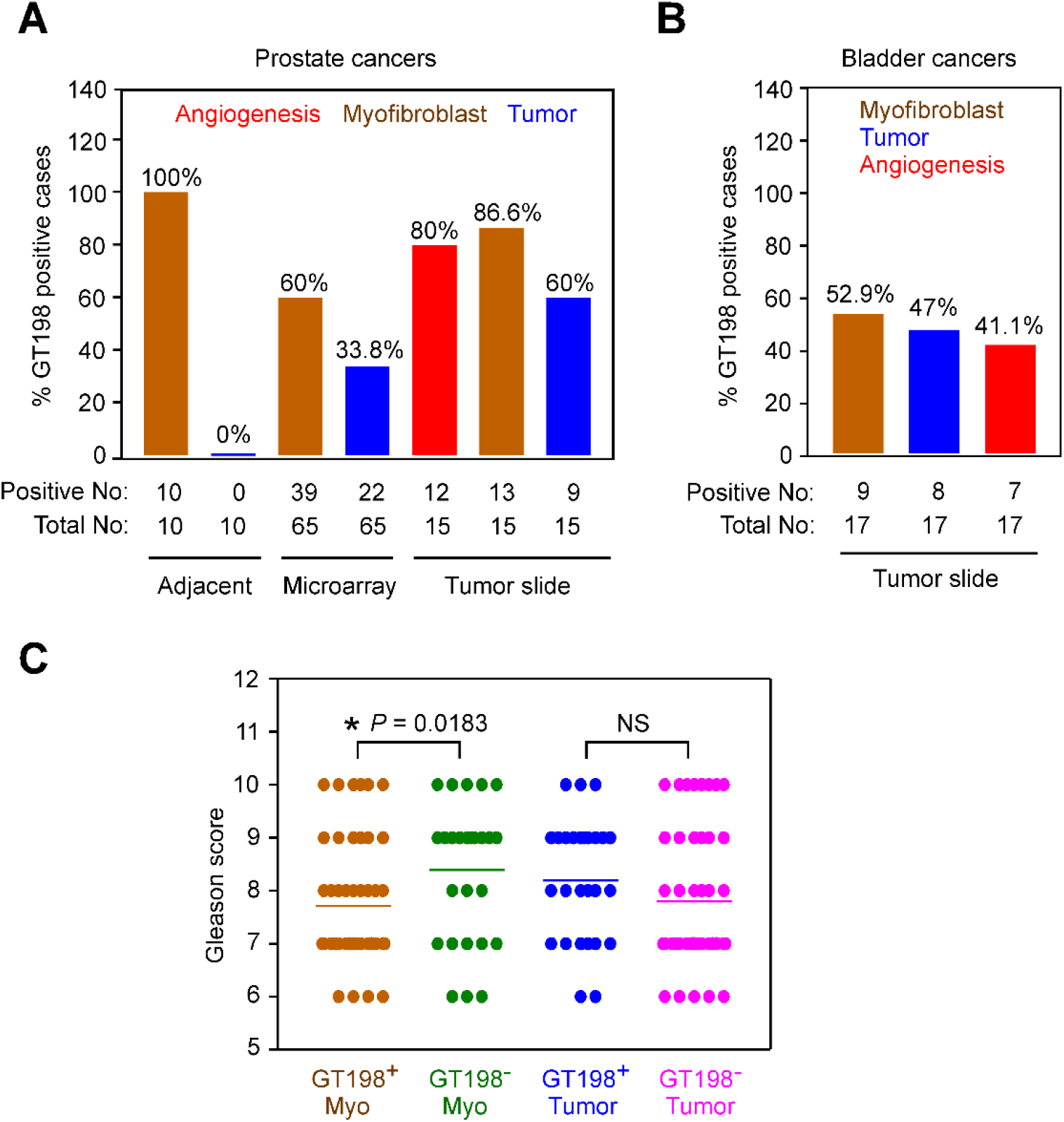
GT198 expression in prostate and bladder cancers. Immunohistochemical staining of GT198 in 65 cases of prostate tumor tissue microarray, 15 cases prostate tumor slides, and 17 cases of bladder tumor slides (Table 1–4). **(A)** Percent of GT198^+^ cases in each type of prostate cancer tissues: pericytes in angiogenic stroma, myofibroblasts, and epithelial tumor cells. **(B)** Percent of GT198^+^ cases in each type of bladder cancer tissues. **(C)** Comparison of Gleason scores in prostate cancers with GT198^+^ or GT198^-^ myofibroblasts and tumor cells. GT198^+^ myofibroblasts have significant association with the cases having lower Gleason scores, which are early stages of cancers. \**P*<0.05. The Gleason sores between GT198^+^ and GT198^-^ tumor cells are insignificantly different. *P*>0.05 (NS).

### GT198^+^ stroma is a predictive biomarker in docetaxel treatment of human bladder cancers

GT198 is expressed in bladder cancers with an overall positive rate of 76.4% (14/17) (**Table 4** and **Figure 3B**). GT198 can be highly expressed in bladder stroma before the evidence of tumor (T0 in cases of B1, B4 and B7) based on the pTNM staging (**Table 4**), although statistical significance cannot be determined due to the small sample size analyzed. B1 and B4 cases also carry *GT198* mutations (**Table 4** and **Figure 5**). The data is consistent with GT198^+^ stroma as very early lesions in bladder cancer (**Figure 2B**).

Clinical information of bladder cancer patients (B1-B17) is shown in **Table 4**. The FFPE slides were immunohistochemical stained with GT198 to determine GT198^+^ myofibroblasts (52.9%, 9/17), GT198^+^ tumor cells (47%, 8/17), and the presence of angiogenesis (41.1%, 7/17) (**Table 4**). The presence of angiogenesis is determined by the presence of blood vessel clusters. Adjacent slides of 15 cases (B1-B15) were Sanger sequencing analyzed for *GT198* mutations with a positive rate of 40% (6/15). Progression-free survival was analyzed by the Kaplan-Meier methods. The results show that GT198^+^ stroma but not GT198^+^ tumor is significantly associated with better progression-free survival, *P*=0.0066, HR=0.1958 (95%CI=0.03-0.08) in docetaxel-treated bladder cancer patients (**Figure 4A-B**). Tumor angiogenesis in stroma is also associated with better progression-free survival, *P*=0.027, HR=0.255 (95%CI=0.06-0.84) (**Figure 4C**). In most cases with GT198^+^ stroma, angiogenesis is present (**Table 4**). This is a first study to suggest that GT198 is a predictive biomarker in docetaxel treatment of bladder cancer. Larger sample size analysis in the future will help to validate the current finding.

**Figure 4.**
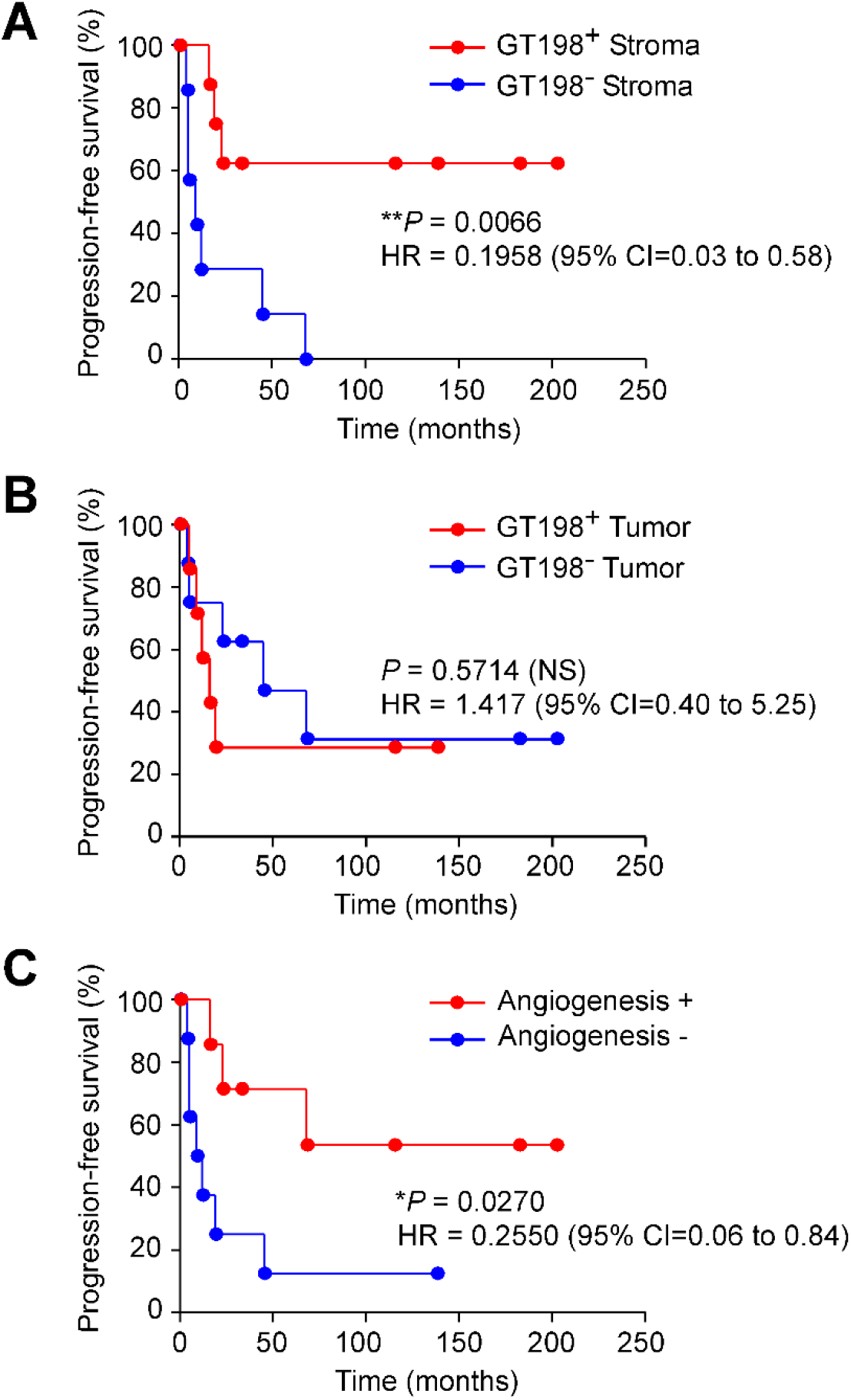
GT198^+^ stroma and angiogenesis are associated with significantly better progression-free survival in docetaxel-treated bladder cancer patients. Clinical information of docetaxel-treated 15 bladder cancer patients is shown in **Table 4**. The 15 FFPE slides were immunohistochemical stained with GT198 for analyzing myofibroblasts and tumor cells. Angiogenesis is determined by the presence of blood vessel clusters. Progression-free survival months and last follow up status are analyzed by the Kaplan-Meier method. **(A)** In docetaxel-treated bladder cancers, GT198 expression in stroma is significantly associated with better progression-free survival. **(B)** GT198 expression status in tumor cells is not associated with the progression-free survival. **(C)** Angiogenesis in tumor stroma, disregarding GT198 expression, is associated with significantly better progression-free survival. *P* values are calculated by log-rank test. \**P* < 0.05. \*\**P*< 0.01. *P*>0.05, not significant (NS).

### GT198^+^ prostate and bladder cancer stroma harbors *GT198* somatic mutations

GT198 expression in stromal blood vessels accompanied by angiogenesis is a strong indicator for the presence of *GT198* somatic mutations in breast, ovarian, and fallopian tube cancers (6–8). Here, we carried out Sanger sequencing analysis of selected prostate and bladder cancers spanning previously identified *GT198* mutation hotspots (**Figure 5**) (7,8). Because stroma proliferation is massive in prostate and bladder cancers, microdissection was not required, and the samples were obtained using GT198-stained adjacent slides to locate the GT198^+^ stroma or tumor.

**Figure 5.**
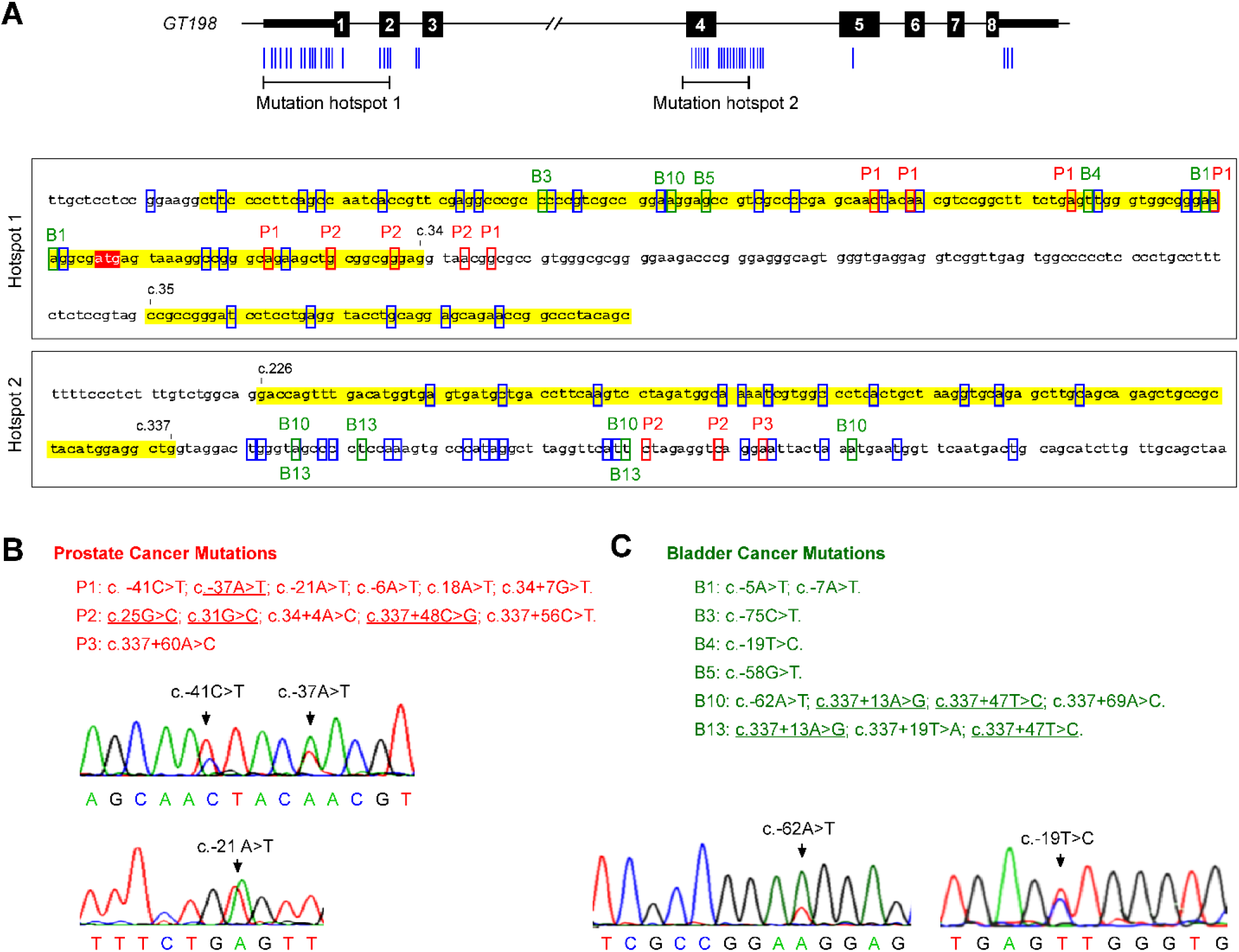
GT198^+^ prostate and bladder cancer stroma harbors *GT198* somatic mutations. **(A)** A diagram of the human *GT198* gene is depicted with numbered boxes as exons and lines as introns. The two mutation hotspots are indicated. Blue bars represent *GT198* germline and somatic mutations from previous studies (6–8,21). Nucleotide sequences of the two mutation hotspots are shown with exons in yellow and start codon in red. Previously identified *GT198* mutations in cancer are indicated in blue boxes. Mutations identified from prostate cancers in this study are indicated in red boxes (P1-P3), and from bladder cancers are indicated in green boxes (B1, B3-B5, B10, B13) (Table 3–4). Mutation numbering is based on the cDNA reference sequence. **(B)** Mutations in prostate cancers with representative sequence trace chromatograms. **(C)** Mutations in bladder cancers with representative sequence trace chromatograms. Nucleotide changes in somatic point mutations are indicated by arrows with wild type sequences shown below. Most mutations are newly identified in this study except that the same mutations found in other cancers from previous studies are underlined.

In four cases of prostate cancer analyzed (**Table 3**), three cases (P1-P3) carry *GT198* somatic mutations with a positive rate of 75% (3/4) (**Figure 5A-B**). Multiple mutations are present in P1 and P2 cases. The P1 prostate cancer stroma carries six *GT198* mutations (**Figure 5B**), has a Gleason score of 6 (**Table 3**), contains mostly GT198^+^ stroma rather than tumor, and has abundant GT198^+^ angiogenic vessels (**Supplemental Figure S1**). This is a typical case that the early stage prostate cancer stroma has multifocal lesions with different *GT198* somatic mutations.

In 15 cases of bladder cancers analyzed, six cases carry *GT198* mutations (40%, 6/15). Mutations can be found in both GT198^+^ stroma and bladder tumors (**Table 4** and **Figure 5C**).

Most *GT198* point mutations identified are newly found in this study, although a few mutations are previously known in other cancers (**Figure 5A**). These mutations are likely deregulating alternative splicing of GT198 as we previously proposed (8), since most of them are not truncating mutations or affecting coding regions (**Figure 5A**).

Because conventional Sanger sequencing requires a mutant cell population carrying the same mutation to yield enough DNA to be detectable, our finding indicates that multifocal and clonally expanded mutant cells are present in prostate and bladder cancers. Together, our results show that high frequency of *GT198* somatic mutations are present in GT198^+^ tumor stroma and tumor cells in human prostate and bladder cancers.

## DISCUSSION

GT198 is an excellent biomarker to reveal the origin of prostate and bladder cancer from their tumor stroma for several reasons. Previous evidence has shown that GT198 is a stem cell regulator affecting blood vessel pericyte stem cells in tumor angiogenesis (33). *GT198* is an oncogene producing mutant tumor stroma driving epithelial cell growth (6,7). GT198 is a nuclear receptor coactivator (4), stimulating hormone-associated target genes which are critical in prostate and bladder cancer development. GT198 is a DNA repair factor (14,19), permitting hypermutation to occur when GT198 itself is functionally defective, similarly to p53. In fact, GT198 and p53 share profound similarities in their structural and functional properties (40–42), and both carry abundant splicing mutations.

A model of GT198 in human prostate and bladder cancer can be proposed in view of historical evidence as well as our current findings (**Figure 6**). Pericytes are stem cells located on capillaries of blood vessels (43), producing vascular smooth muscle cell lineages including myofibroblasts and smooth muscle cells in prostate and bladder. When the *GT198* gene is mutated in pericytes with GT198 overexpression, it causes angiogenesis (33), and produces GT198^+^ stroma (**Figure 1–2**). *GT198* continues to mutate since DNA repair is defective. Hypermutation results in GT198 spicing defects (8), and GT198 protein constitutive activation (7). As a transcriptional coactivator, GT198 may overs timulate steroid hormone receptor-mediated target genes in prostate and bladder, and in turn to induce proliferation of stromal cells and epithelial or urothelial cells (**Figure 6**). This mechanism is likely shared in prostate and bladder cancers. As a biomarker of cancer-initiating stromal cells, GT198 protein may also serve as a drug target (**Figure 4**). This model predicts that GT198 is a common cause and a drug target for both prostate and bladder cancers.

**Figure 6.**
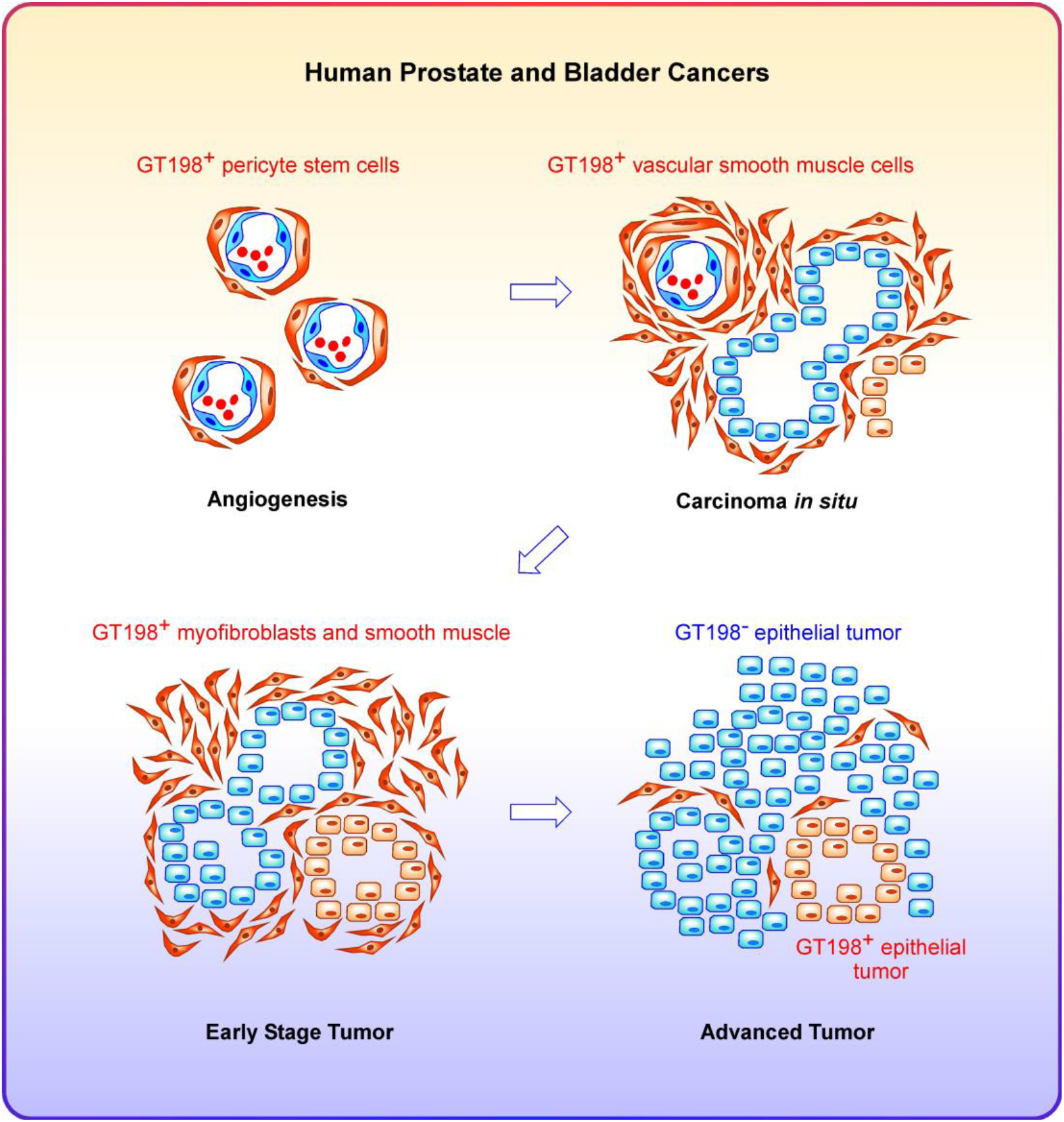
A common origin of human prostate and bladder cancers. A model of oncoprotein GT198 in human prostate and bladder cancer development. Pericytes are stem cells producing vascular smooth muscle cell lineages including blood vessel smooth muscle cells, fibroblasts, myofibroblasts, and smooth muscles in prostate and bladder. When the *GT198* gene is mutated in angiogenic pericytes with overexpression of GT198, GT198^+^ descendent stromal cells create a mutant reactive stroma propelling epithelial cell growth. Epithelial cells can be either GT198^+^ or GT198^-^. At the advanced stages of cancer development, GT198^+^ stromal cells are outgrown by epithelial or urothelial tumor cells. Docetaxel has better treatment efficacy in cancers with GT198^+^ stroma (**Figure 4**), suggesting that it is more effective to target to the origin of cancer, which is shared in human prostate and bladder cancers.

Although high frequency *GT198* mutations are present in prostate and bladder cancers, it is current unclear whether GT198 overexpression is always due to its mutations. Not all GT198^+^ prostate and bladder samples are found mutation positive (**Table 3–4**), however, these could be due to technical reasons. We have only sequenced the two mutation hotspots not the entire *GT198* gene, since DNAs are more fragmented in the FFPE tissues. If a mutation were located outside the hotspots, we would miss to detect. Additionally, scattered mutant stromal cells within outgrowing mutation-negative tumor cells would not be detectable without deep sequencing.

GT198 protein is a predictive biomarker so that it can be a drug target. Paclitaxel was previously found as a direct GT198 inhibitor (39), and its analog docetaxel would inhibit GT198^+^ stromal cells to achieve effective therapy (**Figure 4**). In the future, better GT198 inhibitors can be tested as potential drugs for the treatment of prostate and bladder cancers. As a predictive biomarker, the association of GT198 expression with androgen dependence also needs to be investigated to aid the decision in therapy. In addition, whether *GT198* is responsible for familial prostate cancer can now be analyzed in familial prostate cancer cohorts with segregation by directly sequencing the human *GT198* gene.

In summary, we have found high frequency *GT198* somatic mutations and GT198 oncoprotein overexpression in tumor stroma of human prostate and bladder cancers. GT198^+^ tumor stromal cells include pericyte stem cells, myofibroblasts in prostate cancer, and smooth muscle cells in bladder cancer. Mutant GT198^+^ stromal cells appear at the initial stages of tumor development, resulting in a mutant stroma driving epithelial cell growth. In bladder cancers, the presence of angiogenesis and GT198^+^ stroma are associated with better progression-free survival in docetaxel-treated patients. This implies targeting GT198 at the early stages of cancer is more therapeutic effective. Together, our results suggest the presence of a common origin of both human prostate and bladder cancers, in which mutant GT198^+^ stroma serves as an epicenter of the tumor development. GT198 protein may thus be a therapeutic target in the treatment of human prostate and bladder cancers.

## ACKNOWLEDGEMENTS

We thank Dr. Nahid F. Mivechi for providing reagents. This work was supported in part by the Georgia Cancer Coalition Distinguished Cancer Scholar Award to L.K.

---

**Supplemental Figure S1.**
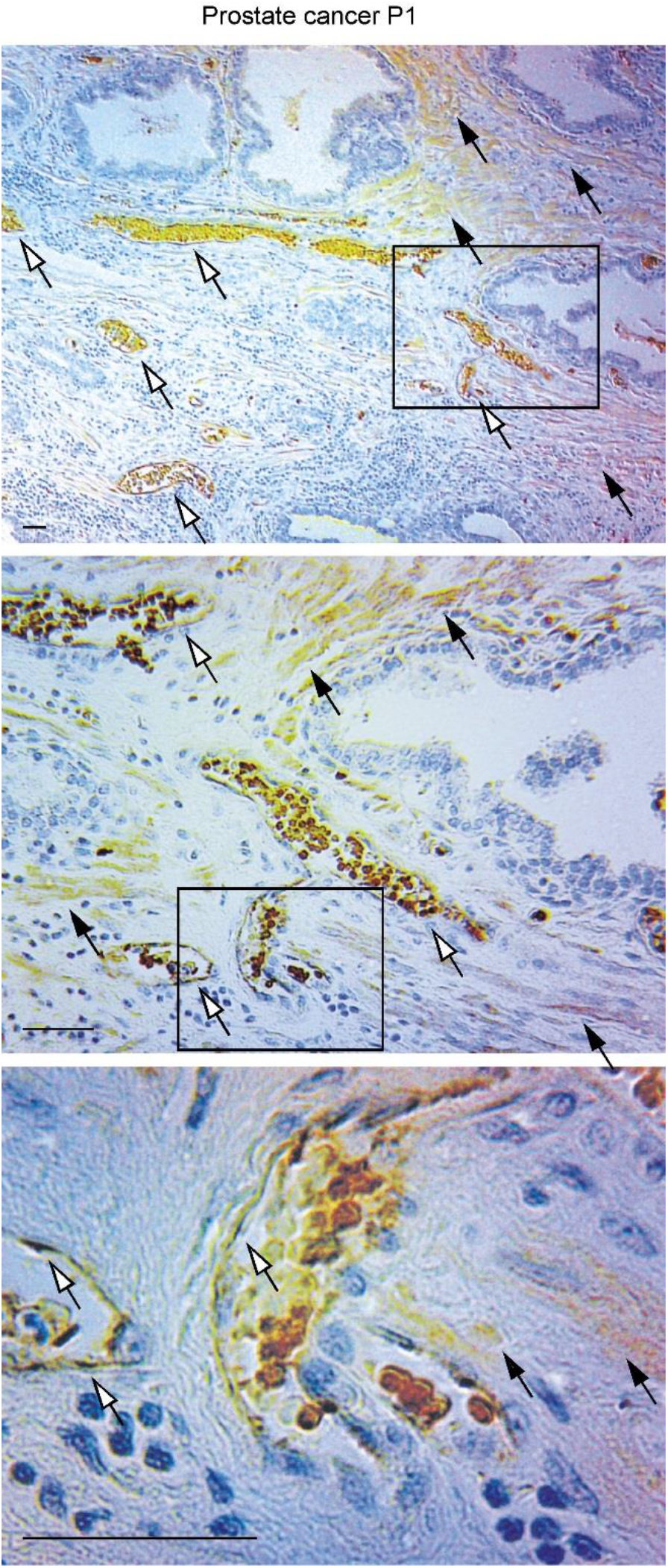
Mutant tumor stroma of prostate cancer P1. Immunohistochemical staining of GT198 in prostate cancer case P1. The adjacent slide was sequencing analyzed to identify six *GT198* somatic mutations (**Figure 5**). Boxed areas are serial enlarged below each panel. White arrows indicate GT198^+^ blood vessels. Black arrows indicate GT198^+^ myofibroblasts. Sections were counter-stained with hematoxylin. Scale bars = 100 μm.

## METHODS

### Study design and human prostate and bladder cancer samples

Institutional Review Board (IRB) approval was obtained following institutional guidelines using de-identified human prostate and bladder cancer paraffin sections. Individual patient consent was not required. Formalin-fixed paraffin-embedded (FFPE) 5-micron sections of human prostate and bladder cancers were obtained from Indiana University School of Medicine, Indiana, IN; Medical College of Georgia, Augusta, GA; and Magee-Womens Hospital, University of Pittsburgh, Pittsburgh, PA. In addition, FFPE tumor tissue microarrays of prostate cancers were from Novus Biologicals, Centennial, CO and US Biomax Inc., Rockville, MD. Pathology diagnosis of all samples was verified through histological examination by pathologists. Prostate and bladder cancer tissue sections were analyzed by immunohistochemistry using anti-GT198. Selected four cases of prostate cancers (P1-P4) and fifteen cases of bladder cancers (B1-B15) were subjected to Sanger DNA sequencing analysis to identify somatic mutations in *GT198* using serial cut adjacent sections.

### Immunohistochemistry

Polyclonal antiserum against GT198 was previously prepared in rabbits (Covance, Denver, PA) (4,7). Affi-gel 10 affinity resins were covalently cross-linked to his-tagged GT198 protein as antigen for affinity purification of anti-GT198 according to the manufacturer’s protocol (Bio-Rad, Hercules, CA). FFPE tumor sections or tumor microarrays were deparaffinized and dehydrated through xylene and ethanol series, followed by antigen retrieval in 10 mM sodium citrate buffer, pH 6.0, containing 0.05% Triton at 90°C for 20 min. Anti-GT198 (1:150) was incubated at 4°C overnight. Antibody binding was detected using biotinylated anti-rabbit secondary antibody followed by detecting reagents (Abcam, Cambridge, MA). Sections were counterstained with hematoxylin. Data was graphed by scattergram using GraphPad Prism software.

### Mutation Analysis

FFPE sections of prostate and bladder tumors were deparaffinized through xylene and 100% ethanol, and air dried. The adjacent sections were first stained by immunohistochemistry to locate the GT198^+^ stroma or tumor, and the positive areas (0.5-1 cm^2^) were removed using a new razor blade and transferred to the tube. Genomic DNA was isolated by DNeasy Tissue Kit reagents (Qiagen, Valencia, CA). DNA samples were PCR amplified and Sanger sequencing analyzed from both directions with repeats in two previously identified mutation hotspot sequences at 5’UTR to exon 2 (c.-109C to c.103A) and at the exon 4-intron 4 junction (c.265C to c.337+116T) (6–8). The entire *GT198* gene was not analyzed due to the fragmentation of DNA in FFPE sections. Primer pairs are:

5’UTR-F: 5’-ggggtcgctttgctcctccggaa-3’,
Intron 1-B: 5’-ctacggagagaaaggcagggga-3’;
Intron 1-F: 5’-cagtgggtgaggaggtcggttga-3’,
Exon 2-B: 5’-agtccgtgttcccgctgtaggt-3’;
Exon 4-F: 5’-gtgagtgatgctgaccttcaagtc-3’,
Intron 4-B: 5’-tacacaaaagccgttagttatcct-3’.

Nucleotide numbering is based on the cDNA reference sequence.

### Kaplan-Meier survival analysis

The study consisted of 15 bladder cancer patients who underwent adjuvant docetaxel treatment at Indiana University, between June 1992 and June 2014. The longest postoperative follow-up duration was 17 years. The primary end point was progression-free survival without bladder tumor reoccurrence from the time of operation to the last day of follow-up. Progression-free survival was estimated by the Kaplan–Meier method. Hazard ratios (HRs) and *P* values were calculated with the GraphPad Prism software using two-sided 0.05 significance level.

### Statistical analysis

Statistical analyses were carried out using GraphPad Prism software. Scattergrams with means are presented using Gleason scores in GT198^+^ cases. *P* values in scattergrams and bar graphs were calculated using unpaired two-tailed t-test. *P* values in Kaplan-Meier curves were calculated by log-rank test. * *P*<0.05, ** *P*<0.01, *** *P* <0.001; NS, not significant. A *P* value of less than 0.05 is considered statistically significant.

